# The *Mstn*^*cmpt-Dl1abc*^ mutation impairs secretion of promyostatin

**DOI:** 10.1101/077412

**Authors:** Viktor Vásárhelyi, Mária Trexler, László Patthy

## Abstract

Hypermuscularity of Compact mouse is caused by a 12-bp deletion in the myostatin gene, but the molecular basis of decreased myostatin activity is unclear since the deletion does not affect the integrity of the growth factor domain. In the present work we show that the deletion causes misfolding and impaired secretion of myostatin precursor with concomitant decrease in myostatin activity. We suggest that some modifier genes that influence the expression of the Compact phenotype of myostatin mutant *MstnCmpt-dl1Abc* mice may exert their action through their role in the Unfolded Protein Response.

## 1. Introduction

Myostatin, a member of the transforming growth factor-β (TGF-β) family of growth factors, is secreted as an inactive precursor, promyostatin [1]. Two molecules of promyostatin are covalently linked via a single disulfide bond present in the C-terminal growth factor domain; the active growth factor myostatin is liberated from promyostatin by proteolytic processing. Furin type proprotein convertases cleave both chains of the precursor at the boundary of the prodomain and the growth factor domain, but the two prodomains and the disulfide-bonded homodimeric growth factor remain associated, forming an inactive complex referred to as a latent complex [2]. Active myostatin is released from this latent complex through degradation of the prodomain by proteases of the bone morphogenetic protein −1 (BMP-1)/Tolloid family of metalloproteinases [3].

Myostatin is a negative regulator of skeletal muscle growth: McPherron *et al.* were the first to show that mice lacking myostatin are characterized by a dramatic increase in skeletal muscle mass [4]. Subsequent studies on several breeds of cattle selected for increased muscle mass have shown that hypermuscularity is caused by an 11-nucleotide deletion in the third exon of the myostatin gene which causes a frameshift that eliminates the myostatin growth factor domain [5, 6, 7]. In the case of a racing-dog breed it was shown that a 2-bp deletion in the third exon of the myostatin gene increases muscle mass and enhances racing performance since the reading frame shift and premature termination removes the major part of the growth factor domain [8]. In the case of a human child it was shown that muscle hypertrophy was caused by a homozygous mutation in the first intron of the myostatin gene that leads to mis-splicing, premature termination and complete loss of myostatin protein [9].

Studies on the hypermuscular Compact mouse, selected for high body weight and protein content, have shown that the Compact phenotype is caused by a 12-bp deletion in the myostatin gene [10]. This deletion, however, did not provide a simple explanation as to how it leads to loss of myostatin activity. Unlike the myostatin mutations mentioned above, this mutation (denoted *Mstn*^*Cmpt-dl1Abc*^) in the prodomain region of the myostatin precursor does not result in premature termination and does not affect the integrity of the growth factor domain [10]. In harmony with the notion that myostatin activity is not zero in homozygous Cmpt/Cmpt mutants, these animals have a less pronounced hypermuscular phenotype than homozygous mice in which the myostatin gene was disrupted by gene targeting [11]. Genetic analyses also revealed that full expression of the hypermuscular Compact phenotype requires the action of modifier loci in addition to *Mstn*^*Cmpt-dl1Abc*^ [12, 13]. These studies also revealed that the modifier loci exerted their effects on muscularity only in the presence of *Mstn*^*Cmpt-dl1Abc*^ [12].

In principle, the *Mstn*^*Cmpt-dl1Abc*^ mutation present in the prodomain region might lead to decreased myostatin activity if it interferes with any of the steps leading to the production of mature myostatin from promyostatin: it might interfere with the folding of the precursor and/or the formation of the covalent homodimer, preventing its secretion or it might impair the liberation of mature myostatin from the precursor and/or the latent complex. In the present work we wished to examine these possibilities, therefore we produced and studied the molecular properties of promyostatins carrying the *Mstn*^*Cmpt-dl1Abc*^ mutation.

Our studies on recombinant proteins expressed in bacterial expression systems have shown that the *Mstn*^*cmpt-Dl1abc*^ deletion causes misfolding of promyostatin. In mammalian expression systems misfolding of *Mstn*^*cmpt-Dl1abc*^ promyostatin impaired its secretion, suggesting that the hypermuscular phenotype of Compact mouse is due to a severe decrease in the level of extracellular myostatin precursor. Based on these findings we suggest that the modifier loci that affect the expressivity of the Compact phenotype [12,13] may encode constituents of the Unfolded Protein Response (UPR) as they may influence the fate of mutant promyostatin.

## 2. Materials and methods

### 2.1. Expression of wild type and *Mstn^Cmpt-dl1Abc^* mutant murine promyostatin in *Escherichia coli*

The cDNA of murine promyostatin was amplifed with the 5′ GGACCGCGGTCAATGAGGGCAGTGAGAGAGAAG 3'sense and 5′ CGGGTCGACTTATCATGAGCACCCACAGCGG 3′ antisense primers from the pCMV6-Entry mouse myostatin plasmid (MR 227629 Origene,) and ligated into the SacII/SalI sites of pPR-IBA2 (IBA) expression vector, yielding the expression plasmid designated as pPR-IBA2_MU_promyostatin.

We used the QuikChange Site-Directed Mutagenesis method to introduce the *Cmpt-dl1Ab* mutation into the coding region of murine promyostatin. The mutagenesis reaction was performed with the 5′ CAAACAGCCTGAATCCAACTTCAAAGCTTTGGATGAGAATGGC 3′ sense and 5′ GCCATTCTCATCCAAAGCTTTGAAGTTGGATTCAGGCTGTTTG 3′ antisense primers and with the pPR-IBA2_MU_promyostatin expression plasmid as template. The mutagenesis deleted the AGGCATTGAAAT nucleotide segment (positions 672-684) of the murine promyostatin cDNA, deleting five amino acids (Leu-Gly-Ile-Glu-Ile, positions 224-228 of murine promyostatin) and inserting a single phenylalanine in the protein.

*E. coli* BL21(DE3) cells were transformed with the pPR-IBA2_MU_promyostatin or pPR-IBA2_MU_ promyostatin_Cmpt_dl1Ab vectors and expression of recombinant proteins was performed with the same method that was used for the production of the recombinant wild type human promyostatin [14].

### 2.2. Expression of *Mstn^Cmpt-dl1Abc^* mutant human promyostatin in *Escherichia coli*

In order to produce the Mstn^Cmpt-dl1Abc^ mutant human promyostatin we used the QuikChange Site-Directed Mutagenesis method to introduce the *Cmpt-dl1Ab* mutation into the coding region of the expression plasmid pPR-IBA2_HU_promyostatin containing the cDNA of human promyostatin [14]. Mutagenesis was performed with the 5′ CAAACAACCTGAATCCAACTTCAAAGCTTTAGATGAGAATGGTC 3′ sense and 5′ GACCATTCTCATCTAAAGCTTTGAAGTTGGATTCAGGTTGTTTG 3′ antisense primers. As in the case of the *Mstn*^*Cmpt-dl1Abc*^ mutant of murine promyostatin, the mutation replaced residues Leu-Gly-Ile-Glu-Ile (positions 223-227 of human promyostatin) by a phenylalanine residue.

*Escherichia coli* BL21(DE3) cells were transformed with the pPR-IBA2_HU_Promyostatin_Cmpt_dl1Ab plasmid and expression, purification and refolding of recombinant mutant promyostatin was performed with the protocol used for the production of the recombinant wild type human promyostatin [14].

### 2.3. Expression of wild type and *Mstn^Cmpt-dl1Abc^* mutant promyostatins in HEK 293 human embryonic kidney cells

The cDNAs of wild type mouse or human promyostatin were cloned into the pENTRY-IBA51 entry vector (IBA) and the resulting pENTRY-IBA_MU_Promyostatin and pENTRY-IBA_HU_Promyostatin plasmids were used as templates of the Quickchange mutagenesis reaction; mutagenesis was performed with the primers given in sections 2.2 and 2.3. After sequence verification the inserts encoding the wild type and mutant promyostatins were transferred to the pCSG-IBA142 mammalian expression vector.

The HEK 293 cells used to express promyostatins were cultured in DMEM medium containing 10% heat inactivated fetal bovine serum in the presence of 5% CO_2_ at 37°C. For transfection 3 ml aliquots containing 5 × 10^5^ HEK 293 cells were seeded in each well of a 6-well plate and the cells were allowed to attach for 24 h. The cells were transfected with 3-3 µg of pCSG-IBA142-HU_Promyostatin, pCSG-IBA142-HU_Promyostatin*_*Cmpt-dl1Ab, pCSG-IBA142-MU_Promyostatin, pCSG-IBA142-MU_Promyostatin*_*Cmpt-dl1Ab or pCSG-IBA142-control plasmids with Fugene HD transfection reagent (Promega) according to the recommendations of the manufacturer. 24 h after transfection the culture media were changed to serum-free DMEM and transient protein expression was allowed for 48 h.

Culture fluids were concentrated to 200 µl, cells were lysed with 200 µl icecold RIPA buffer (50 mM TRIS, 150 mM NaCl, 1% Triton X-100, 0.5% Na-deoxycholate, 0.1% SDS pH 8.0) containing proteinase inhibitors (2 μg/ml Aprotinin, 5μg/ml Leupeptin, 1 mM PMSF, 5 mM EDTA, 1 mM EGTA) and 15 µl aliquots were subjected to SDS-PAGE on 12 % gels. Western blot analysis of the gels was performed using antibody AF788, specific for the growth factor region of promyostatin (R&D Systems). GAPDH (glyceraldehyde-3-phosphate dehydrogenase) used as loading control was visualized with Anti-GAPDH mouse mAb 6C5 (Calbiochem).

### 2.1. Bioinformatics

Secondary structure of wild type and mutant mouse and human promyostatin were predicted with Jpred3 (http://www.compbio.dundee.ac.uk/jpred) that provides a three-state (α-helix, β-strand and coil) prediction of secondary structure at an accuracy of 81.5% [15].

The homology models of murine and human promyostatin were generated with the automated protein structure homology-modelling server SWISS-MODEL Version 8.05 (http://swissmodel.expasy.org/ [16, 17]. The template was identified from the SWISS-MODEL template library (SMTL version 2014-07-09, PDB release 2014-07-03) with the automated template search of the pipeline. The models were based on the selected 3rjr.1.A file that contains the structure of porcine proTGFbeta1 [18]. The sequence identities of murine and human myostatin with porcine proTGFbeta1 were 31.53 % and 31.68 %, respectively over the aligned regions. The models generated by the pipeline were analyzed with Swiss-PdbViewer DeepView version 4.1 (http://spdbv.vital-it.ch/ [19]. For further details of homology modeling see the online Supplementary files 1 and 2 at http://journals.cambridge.org/grh.

The list of genes present in modifier loci that affect expression of the hypermuscular Compact phenotype of mice carrying Mstn^Cmpt-dl1Abc^ [12,13] were identified using the Mouse Genome Informatics database (http://www.informatics.jax.org) and their possible effect on the Compact phenotype was evaluated based on the functional annotation of the proteins in the Swiss-Prot database (http://www.uniprot.org/uniprot/).

## 3. Results

### 3.1. The Mstn^cmpt-Dl1abc^ mutation impairs the folding of promyostatin expressed in Escherichia coli

Wild type and *Mstn*^*cmpt-Dl1abc*^ mutant promyostatins were expressed in *Escherichia coli* BL21(DE3) according to the protocol described previously for wild type human promyostatin [14]. Although this protocol yielded biologically active, homodimeric proteins in the case of wild type [14] and K153R mutant human promyostatin [20], in the case of murine myostatin, neither the wild type nor the *Mstn*^*cmpt-Dl1abc*^ mutant protein were expressed. Since the cDNAs of murine and human promyostatin are only 91.8 % identical at the nucleotide sequence level, it seems possible that the codon usage of murine promyostatin cDNA is less optimal for expression in *Escherichia coli,* than that of the human protein.

To overcome this problem we have introduced the *Mstn*^*cmpt-Dl1abc*^ mutation into human promyostatin. Human promyostatin carrying the *Mstn*^*cmpt-Dl1abc*^ mutation was similar to the wild type protein in as much as it was also expressed in *Escherichia coli* BL21(DE3) and also accumulated in inclusion bodies. In the case of the mutant protein, however, the refolding protocol yielded covalent polymers (Fig. 1), indicating that the deletion has deleterious effects on the structure, folding and homodimerization of promyostatin.

**Figure 1.**
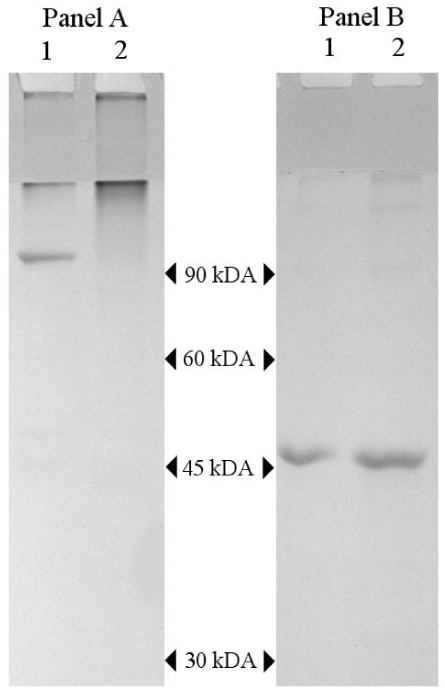
The *Mstn*^*cmpt-Dl1abc*^ mutation impairs the folding of human promyostatin. Wild type and mutant promyostatins were expressed in *Escherichia coli* and refolded according to the protocol described earlier [14]. In the case of wild type promyostatin (lane 1) the majority of the recombinant proteins formed homodimeric promyostatin, but in the case of promyostatin carrying the *Mstn*^*cmpt-Dl1abc*^ mutation (lane 2) the majority of the proteins formed covalent polymers. Non-reduced (-DTT, **panel A**) and reduced samples (+DTT, **panel B**) were run on 12 % SDS-PAGE and were visualized by staining with Coomassie Brilliant Blue R-250. The numbers indicate the Mr values of proteins of the Low Molecular Weight Calibration Kit.

### 3.2. Effect of the *Mstn^cmpt-Dl1abc^* mutation on the structure of promyostatin

Our observation that the *Mstn*^*cmpt-Dl1abc*^ mutation causes misfolding of promyostatin suggested that the mutation might affect a region that is critical for the structural integrity, folding and homodimerization of this protein. Although, the three-dimensional structure of promyostatin is unknown, its homology with pro-TGF-β1 suggested that pro-TGF-β1 and promyostatin have similar secondary structural elements in equivalent positions and that the spatial arrangement of these elements is also similar [18]. In harmony with this suggestion secondary structure prediction with Jpred 3 predicts similar secondary structural elements in equivalent positions with high confidence. Furthermore, homology modeling of human and murine promyostatin using the crystal structure of pro-TGF-β1 as template (Table 1.) confirmed the suggestion that the topology of promyostatin is similar to that of pro-TGF-β1 (see Supplementary file). Although the overall model quality (Global Model Quality Estimation score) of the homology models of human and murine promyostatin is limited by their moderate (<40%) sequence similarity with the template, the Local Quality Estimates of the conserved secondary structural elements are very high since their local sequence similarities are >80% (see Supplementary file).

**Table 1:** Summary of homology modeling of human and mouse promyostatin. The homology models of murine and human promyostatin were generated with the automated protein structure homology-modelling server SWISS-MODEL Version 8.05.

|  |  |
| --- | --- |
| **Template protein:** | transforming growth factor beta-1 |
| Template structure: | X-ray |
| PDB ID of template structure: | 3rjr.1.A |
| Resolution of structure: | 3.05Å |
| **Target protein 1:** | human promyostatin |
| Sequence similarity with template: | 0.34 |
| Range of similarity (residues): | 36 – 375 |
| Coverage: | 0.86 |
| GMQE score*: | 0.55 |
| **Target protein 2:** | murine promyostatin |
| Sequence similarity with template: | 0.34 |
| Range of similarity (residues): | 37 – 376 |
| Coverage: | 0.85 |
| GMQE score*: | 0.55 |

\*GMQE scores (Global Model Quality Estimation score, ranging between 0-1) combine properties from the target-template alignment and reflect the expected accuracy of a model built with that alignment and template. Higher GMQE scores indicate higher reliability.

In the crystal structure of homodimeric pro-TGF-β1 [18] (and the homology models of promyostatins), the N-terminal parts of the two prodomains, consisting of the α1 and α2 helices, provide the ‘straitjacket’ that encircles and shields each growth-factor monomer of the homodimer. The C-terminal parts of the two prodomains (that provide the ‘arm domains’ that connect the two prodomains in the homodimeric precursor) contain three distinct structural regions. A four-stranded β-sheet (consisting of β1, β3, β6 and β10 strands and buried by the α2, α3 and α4 helices) is in close proximity of the growth factor monomer of the same polypeptide chain, whereas a four-stranded β-sheet (consisting of β2, β4, β5 and β7 strands) and β-strands β8 and β9 extend to link the two arm domains of the homodimeric precursor in a ‘bowtie’ [18].

The *Mstn*^*cmpt-Dl1abc*^ mutation that deletes 12 nucleotides of the murine myostatin gene, results in the replacement of residues Leu-224, Gly-225, Ile-226, Glu-227 and Ile-228 by a phenylalanine residue [10]. Since residues 224-228 correspond to the five residues that form the β7 strand of the arm domain (Fig. 2) it is plausible to assume that deletion of this β7 strand may affect the formation of the four-stranded β-sheet that appears to be critical for the folding and homodimerization of promyostatin. It should be noted that secondary structure prediction with Jpred 3 predicts with high confidence the presence of the β7 strand in wild type murine and human promyostatin, but the β7 strand is missing from predictions for murine and human promyostatins carrying the *Mstn*^*cmpt-Dl1abc*^ mutation.

**Figure 2.**
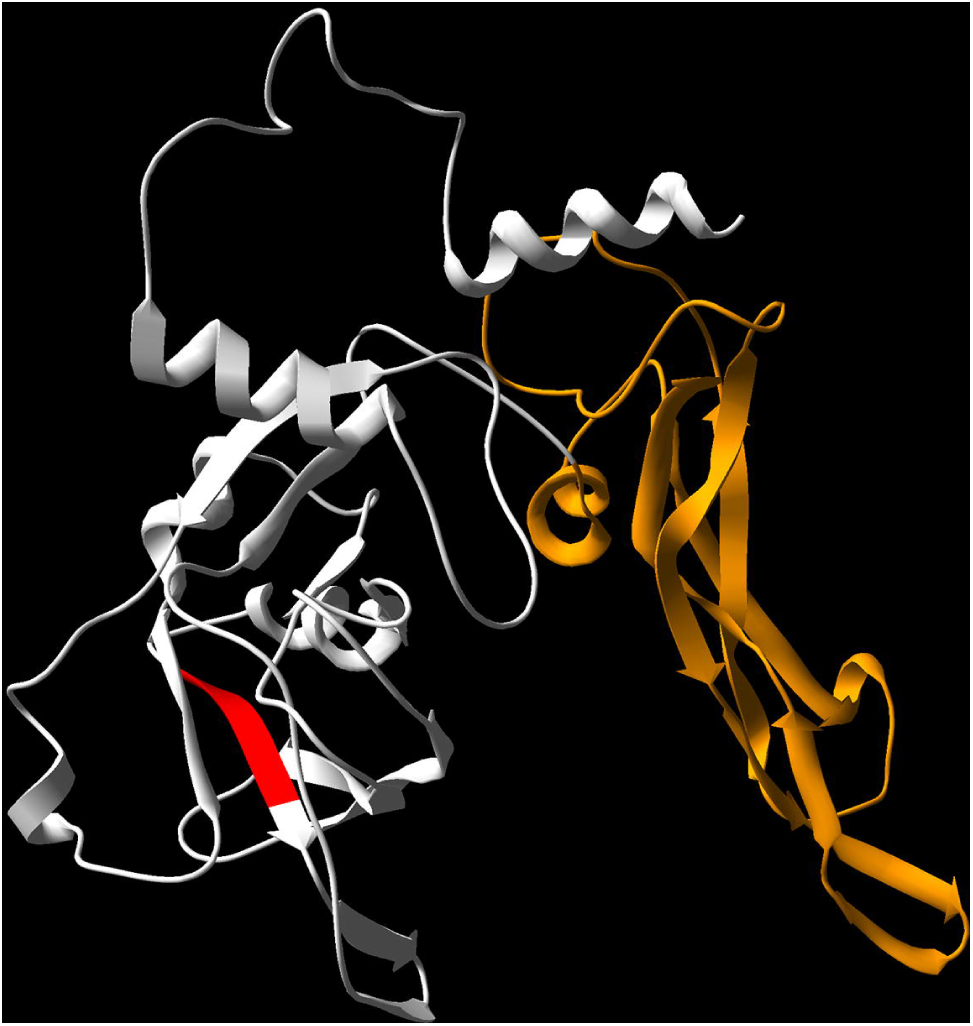
Position of the residues deleted by the *Mstn*^*cmpt-Dl1abc*^ mutation in the homology model of mouse promyostatin. The figure shows the homology model of murine promyostatin highlighting the positions of the residues deleted in the Compact mutant (residues in red). Note that the Compact mutation deletes the β7 strand of the prodomain of promyostatin. The growth factor domain of promyostatin is shown in gold. The homology model was computed using the structure of pro-TGF-1 as a template.

Our observation that the *Mstn*^*cmpt-Dl1abc*^ deletion interferes with the folding and homodimerization of promyostatin is in harmony with the suggestion that the β7 strand is critical for the integrity of the four-stranded β-sheet formed by β2, β4, β5 and β7 of promyostatin. Nevertheless, the failure of *Mstn*^*cmpt-Dl1abc*^ mutant promyostatin to attain a native-like structure *in vitro* does not exclude the possibility that in mammalian cells *Mstn*^*cmpt-Dl1abc*^ mutant promyostatin may acquire native-like structure. To check this possibility we have studied the expression of *Mstn*^*cmpt-Dl1abc*^ mutant promyostatins in mammalian cells.

### 3.3. The *Mstn^cmpt-Dl1abc^* mutation impairs secretion of promyostatin in HEK 293 cells

In our earlier work we have shown that human HEK 293 cells transfected with wild type or K153R mutant promyostatin secrete the precursor efficiently [20]. In harmony with these data, the culture fluids of HEK 293 cells transfected with wild type mouse or human promyostatin contained significant amounts of secreted promyostatin. However, the culture fluids of cells transfected with mouse and human promyostatin carrying the *Mstn*^*cmpt-Dl1abc*^ mutation did not contain detectable amounts of promyostatin (Fig. 3).

**Figure 3.**
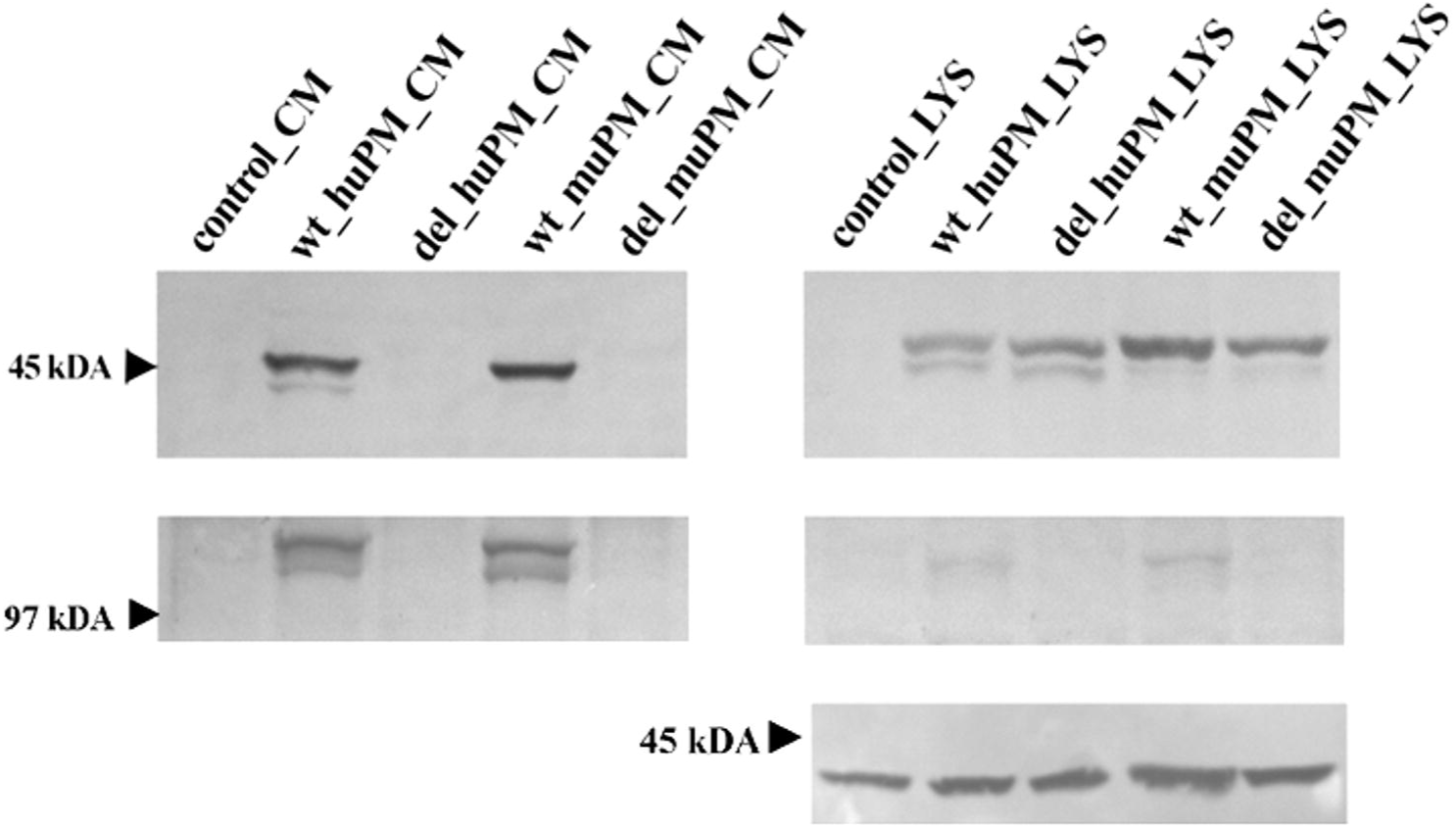
The *Mstn*^*cmpt-Dl1abc*^ mutation impairs secretion of promyostatin in HEK 293 cells. HEK 293 human embryonic kidney cells were transfected with pCSG-IBA142 mammalian expression vector (control), or pCSG-IBA142 vectors harboring the cDNA of wild type human promyostatin (wt huPM), wild type mouse promyostatin (wt muPM), human promyostatin carrying the *Mstn*^*cmpt-Dl1abc*^ deletion (del huPM) or mouse promyostatin carrying the *Mstn*^*cmpt-Dl1abc*^ deletion (del muPM). Transient protein expression was allowed for 48 h in serum-free DMEM then reduced samples of the culture media (CM) and cell lysates (LYS) were subjected to SDS-PAGE on 12% gels. Promyostatin was visualized on Western blots using antibody AF788, specific for the growth factor region of promyostatin. The left hand panels show the results of the analysis of the culture media and the right hand panels show the results obtained by analysis of cell lysates. The arrows indicate the positions of ovalbumin (45 kDa) and phosphorylase b (97 kDa) of the Low Molecular Weight Calibration Kit. Note that wild type human and mouse promyostatin are secreted into the culture medium (wt huPM_CM and wt muPM_CM), but promyostatin could not be detected in culture fluids of control cells (control_CM) or cells transfected with promyostatins carrying the *Mstn*^*cmpt-Dl1abc*^ deletion (del huPM_CM and del muPM_CM). Also note that promyostatin is present in comparable amounts in the lysates of cells transfected with wild type or mutant promyostatin (cf. wt huPM_LYS vs. del huPM_LYS and wt muPM_LYS vs. del muPM_LYS). Top row: reduced samples; lower row: non-reduced samples. In the case of cell lysates GAPDH was used as a loading control (bottom panel).

Analyses of reduced samples of lysates of HEK 293 cells revealed that cells transfected with wild type and *Mstn*^*Cmpt-Dl1abc*^ mutant promyostatins were similar in as much as they contained comparable amounts of intracellular promyostatin (Fig. 3). Analyses of non-reduced samples have shown that lysates of HEK 293 cells transfected with wild type promyostatins contained promyostatin dimers. Promyostatin dimers were, however, missing from lysates of cells expressing mutant promyostatins, indicating that the mutant proteins failed to form disulfide-bonded homodimers.

## 4. Discussion

In the present work we have shown that the *Mstn*^*cmpt-Dl1abc*^ mutation that deletes 12 nucleotides of the myostatin gene causes misfolding and impaired secretion of myostatin precursor with concomitant decrease in myostatin activity.

Our observation that lack of secretion of *Mstn*^*Cmpt-Dl1abc*^ mutant promyostatins did not lead to a significant increase in the amount of intracellular promyostatin (as compared with wild type promyostatin) was somewhat surprising since there is a general consensus that mutations that affect secretory signals usually lead to intracellular accumulation of the mutant proteins (e.g. [21, 22, 23]). The most plausible explanation for the lack of significant accumulation of mutant promyostatin is that terminally misfolded, improperly assembled *Mstn*^*Cmpt-Dl1abc*^ mutant promyostatins are recognized by the quality-control system of the endoplasmic reticulum and are degraded by the proteasome [24].

In our earlier work we hypothesized that, although the Mstn^cmpt-Dl1abc^ mutation may seriously decrease myostatin activity, this activity is not zero in homozygous Cmpt/Cmpt mutants, therefore modifier genes may have a significant influence on the Compact phenotype [10]. The search for modifier genes affecting the expression of hypermuscularity in the Compact mouse was based on the assumption that the Compact line, in addition to achieving homozygosity for the *Mstn*^*Cmpt-dl1Abc*^ mutation, also accumulated modifier alleles that increased the expression of hypermuscularity in the Compact mouse. These studies have revealed significant association with hypermuscularity for markers on chromosomes 1, 3, 5, 7, 11, 16, and X, the strongest association was found for markers on chromosomes 16 and X [12,13].

In principle, modifiers of the Compact phenotype may enhance hypermuscularity by increasing the penetrance of the *Mstn*^*Cmpt-dl1Abc*^ mutation, through their influence on myostatin activity, their effect on downstream components of the myostatin signaling pathway or their effect on other genes regulating muscle development.

Earlier surveys of the genes present in the chromosomal regions showing strong association with hypermuscularity of Compact mice identified a number of candidate genes that might play some of these roles in the regulation of muscle mass [12,13] see Table 2. For example, it was pointed out that regions containing modifier loci on chromosome 7 contain gene MyoD1 of Myoblast determination protein 1, a key regulator of skeletal myogenesis, as well as gene Pcsk6 of proprotein convertase subtilisin/kexin type 6, an enzyme involved in the proteolytic processing of precursors of TGF-βfamily members. Similarly, it was noted that a region of chromosome 16 associated with hypermuscularity of Compact mouse contains gene Chrd of chordin, a protein known to bind members of the TGF-βfamily and sequester them in latent complexes. In the case of chromosome 1 a modifier of the expressivity of the Compact phenotype was mapped to the same region as Myog of myogenin, a downstream target of myostatin [13]. Along the same lines, it was emphasized that the chromosomal region corresponding to the strongest peak on chromosome X [12] contains gene Ar of androgen receptor and it was suggested that Ar might act as a modifier of the Compact phenotype through its influence on the expression of myostatin.

**Table 2:**
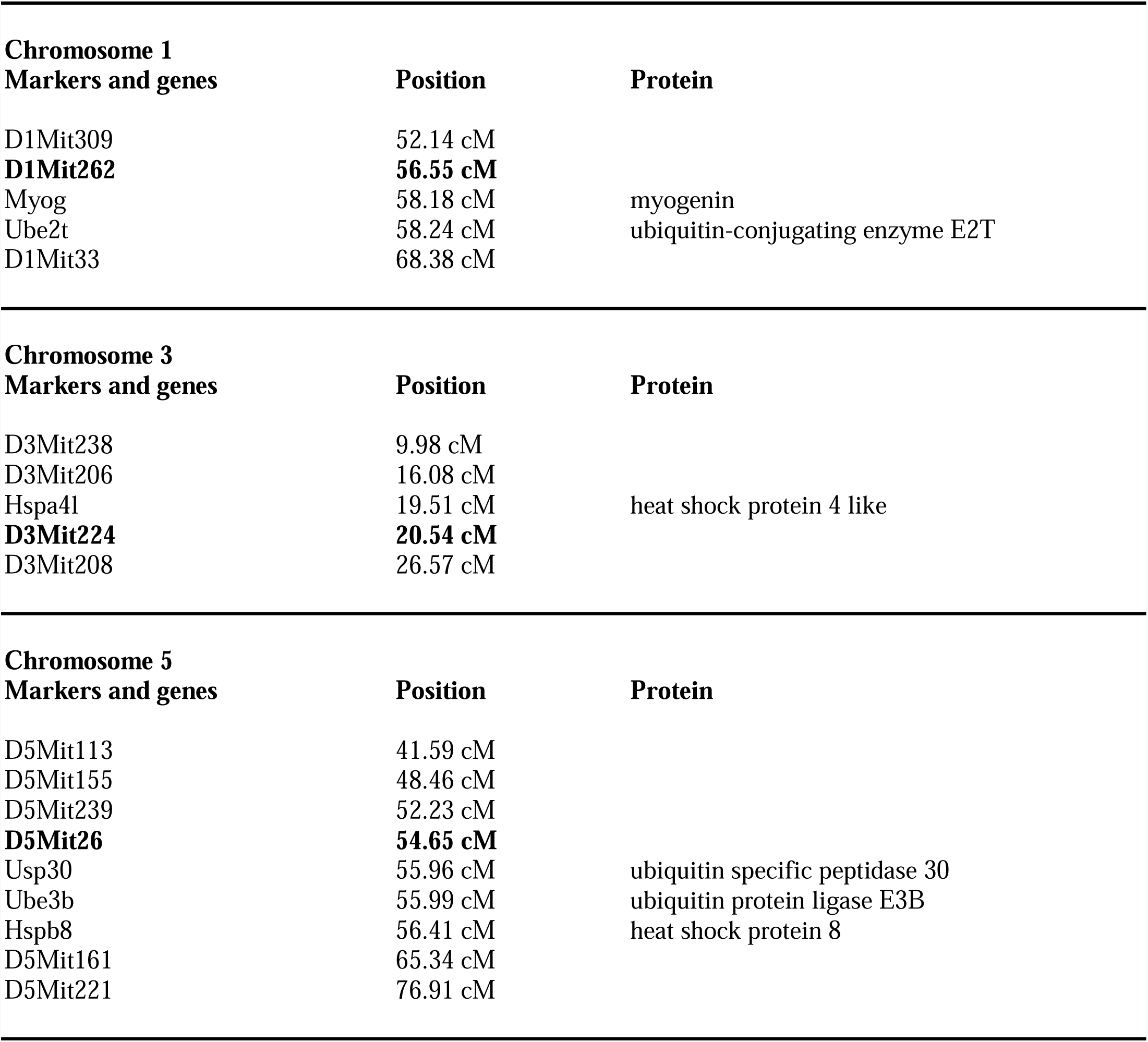
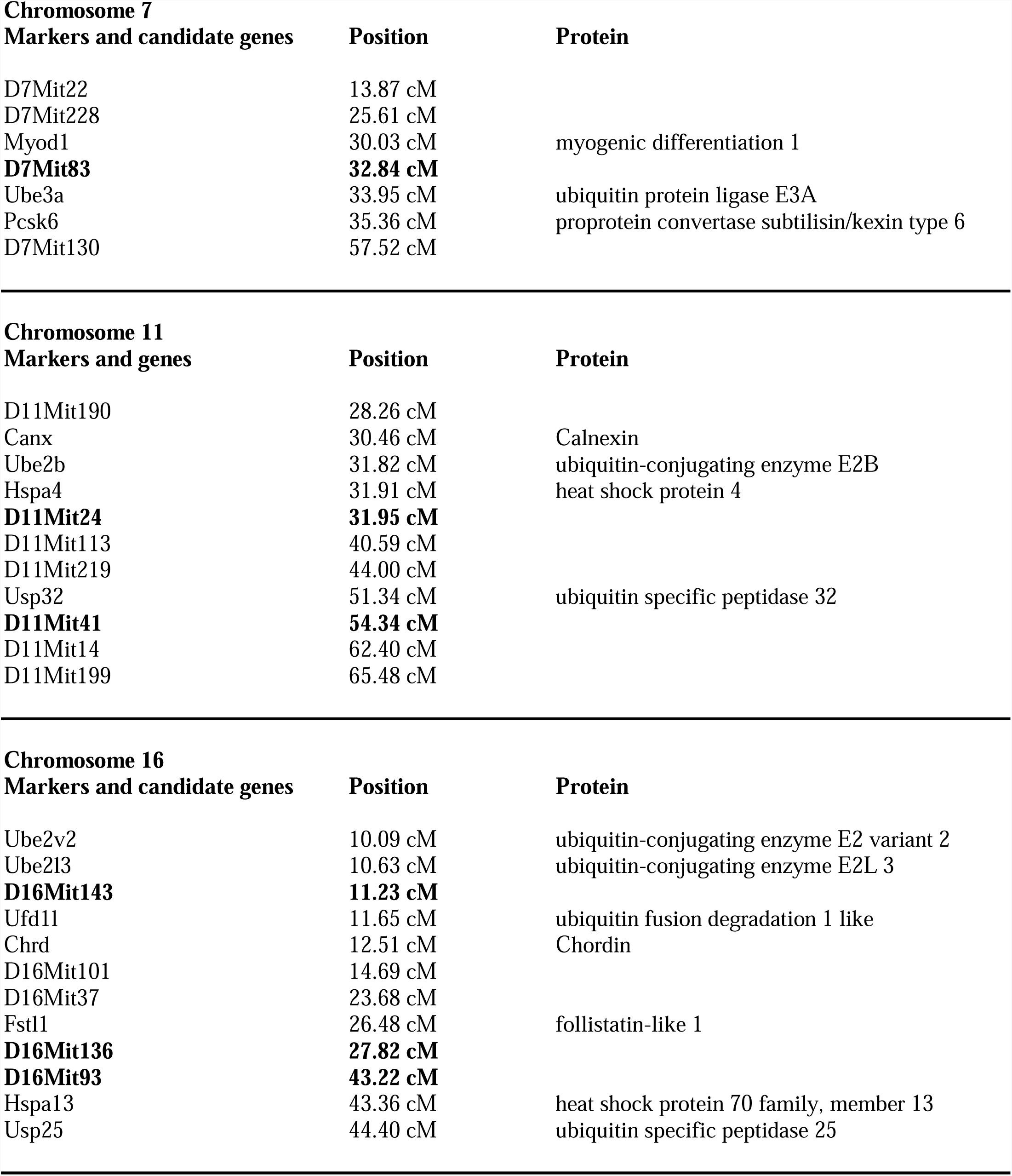
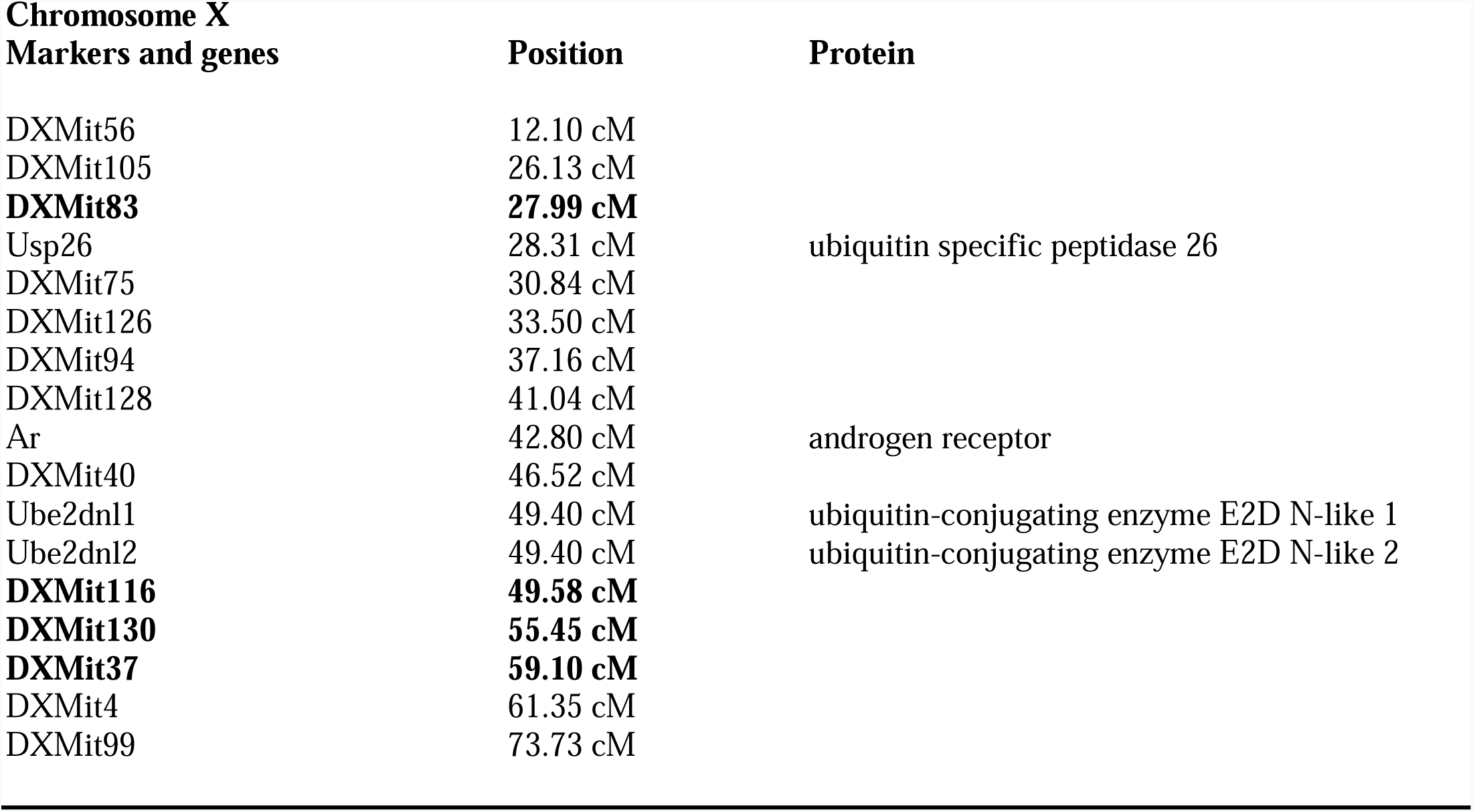
Markers and selected genes near major modifier loci influencing the compact phenotype in the myostatin mutant (*Mstn*^*cmpt-Dl1abc*)^ compact mouse. For each chromosome the markers closely linked to major modifier loci are in bold.

It must be emphasized, however, that the studies that identified regions on Chromosomes 1, 3, 5, 7, 11, 16 and X showing strong association with the hyper-muscularity of compact mice were of relatively low resolution, the association of modifier loci with selected markers showed rather broad (several cM wide) peaks (see [10,12,13]). The putative modifier genes mentioned above are usually separated by several cM from the markers showing strongest association with the modifier loci (Table 2). The problems caused by low resolution modifier-maps may be illustrated by gene Ar of androgen receptor that was suggested to act as a modifier of the Compact phenotype as it is found in the chromosomal region corresponding to the strongest peak on chromosome X [12]. More recent studies refuted this assumption: quantitative analyses of gene expression did not show differences between mRNA levels of androgen receptor in Compact and control mice, indicating that the androgen receptor gene is not the true X-linked modifier of the Compact phenotype [25,26]. Furthermore, finer mapping of the modifier loci on chromosome X has succeeded in narrowing down the modifiers to two regions that are distinct from the region containing the gene of androgen receptor [26].

In view of our finding that the *Mstn*^*Cmpt-dl1Abc*^ mutation leads to misfolding and impaired secretion of mutant promyostatin, we think that more attention should be paid to genes of proteins involved in the quality control system of the endoplasmic reticulum. We assume that some mutations affecting constituents of Unfolded Protein Response system may enhance the expressivity of the *Mstn*^*Cmpt-dl1Ab*^^c^ mutation if they increase the efficiency of the removal of *Mstn*^*Cmpt-dl1Ab*^^c^ mutant promyostatin. In other words, mutations are expected to increase hypermuscularity of Compact mouse if they decrease the activity (decrease expression and or decrease molecular activity) of proteins assisting protein folding (e.g. chaperones) of the *Mstn*^*Cmpt-dl1Ab*c^ mutant promyostatin or increase the activity (increase expression) of proteins assisting removal of the misfolded protein (e.g. constituents of the ubiquitin-dependent protein degradation system). Our survey of the modifier regions of chromosomes 1, 3, 5, 7, 11, 16, and X that show strong association with hypermuscularity of Compact mice [12, 13, 26] has indeed identified a number of genes that might affect the folding and rescue or degradation of *Mstn*^*Cmpt-dl1Ab*c^ mutant promyostatin (Table 2). It should be noted, however, that the extended chromosomal regions that are suspected to contain the modifier loci encode hundreds of genes, only a small fraction of which have a role in protein quality control.

Considering modifier loci on chromosomes 1, 3, 5, 7, 11, 16, and X individually, on chromosome 1 there is a clear maximum between the markers D1Mit309 and D1Mit33, close to D1Mit262 (Table 2). As pointed out earlier [13], this region contains the gene Myog for myogenin, however, in the vicinity of D1Mit262 (56.55 cM) and Myog (58.18 cM) we also find gene Ube2t (58.24 cM) of ubiquitin-conjugating enzyme E2 T (UniProt: Q9CQ37) that catalyzes covalent attachment of ubiquitin to other proteins, marking them for degradation.

On chromosome 3 a narrow region of strong effect was found on the segment defined by markers D3Mit206 and D3Mit224, close to D3Mit224 [12]. In the vicinity of D3Mit224 (20.54 cM) we find gene Hspa4l (at 19.51 cM) of heat shock protein 4 like (UniProt: P48722), a chaperone that aids protein folding.

On chromosome 5 a narrow region of strong effect was defined by markers D5Mit155, D5Mit239, D5Mit26 and D5Mit161, close to D5Mit26 [12]. In the vicinity of D5Mit26 (54.65 cM) we find gene Usp30 (55.96 cM) for ubiquitin specific peptidase 30 (UniProt: Q3UN04) and gene Ube3b (55.99 cM) for ubiquitin protein ligase E3B (UniProt: Q9ES34), two proteins involved in ubiquitin-dependent degradation of proteins. This region also contains gene Hspb8 (56.41 cM) for heat shock protein 8 (UniProt: Q9JK92), a protein with chaperone activity.

On chromosome 7 a modifier region was mapped close to D7Mit83 [12]. As noted previously [12], genes of Myod1 and Pcsk6 map to this region. However, we note that gene Ube3a (33.95 cM) of ubiquitin protein ligase E3A (UniProt: O08759) is even closer to D7Mit83 (32.84 cM) than Myod1 (30.03 cM) or Pcsk6 (35.36 cM). It should be pointed out that ubiquitin protein ligase E3A functions as a cellular quality control ubiquitin ligase by helping the degradation of misfolded proteins.

Broad regions of effect were found on chromosome 11 with two major peaks at D11Mit24 and D11Mit41 [12]. Close to D11Mit24 (31.95 cM) we find gene Hspa4 (31.91 cM) of heat shock 70 kDa protein 4 (UniProt: Q61316), a constituent of the Unfolded Protein Response that is involved in chaperone-mediated protein complex assembly. The same region also contains gene Canx (30.46 cM) of calnexin (UniProt: P35564), a protein that plays a major role in the quality control apparatus of the endoplasmic reticulum by the retention of incorrectly folded proteins, as well as Ube2b (31.82 cM) of ubiquitin-conjugating enzyme E2B (UniProt: P63147). A possible candidate gene in the peak region defined by D11Mit41 (54.34 cM) is gene Usp32 (at 51.34 cM) of ubiquitin specific peptidase 32 (Uniprot: F8VPZ3), an enzyme involved in ubiquitin-dependent protein degradation.

Chromosome 16 also has an extended modifier region with three peaks at D16Mit143, D16Mit136 and D16Mit93 [12]. In the vicinity of D16Mit143 (11.23 cM) we find gene Ube2l3 (10.63 cM) of ubiquitin-conjugating enzyme E2L3 (UniProt: P68037) that is involved in the selective degradation of short-lived and abnormal proteins, as well as Ufd1l (11.65 cM) of ubiquitin fusion degradation 1 like (UniProt: P70362), an essential component of the complex that exports misfolded proteins from the endoplasmic reticulum to the cytoplasm, where they are degraded by the proteasome. Note that these genes are closer to the D16Mit143 marker (11.23 cM) than Chordin (12.51 cM) that was suggested earlier as a potential modifier gene.

In the sharp peak defined by the D16Mit136 marker (27.82 cM) we find gene Fstl1 (26.48 cM) of follistatin-like 1 (Q62356) that, similarly to other follistatin-related proteins, is an antagonist of growth factors of the TGF-ΰfamily [27, 28]. Close to the peak at D16Mit93 (43.22 cM) we find gene Hspa13 (43.36 cM) of the chaperone, heat shock protein 70 family, member 13 (Q8BM72) and gene Usp25 (44.40 cM) of ubiquitin specific peptidase 25 (P57080), a constituent of the Ubiquitin Proteasome System.

Chromosome X showed a broad and exceptionally strong region of effect extending from DXMit126 to DXMit99, with a single peak at DXMit128 [12]. Although the gene of androgen receptor present in this region appeared to be an attractive candidate for the X-linked modifier of the Compact phenotype, subsequent studies refuted this assumption: 1. Neither the structure nor the level of expression of androgen receptor were different in Compact and control mice; 2. Finer mapping of the modifier loci on chromosome X excluded the androgen receptor gene as the major X-linked modifier of the Compact phenotype [25, 26]. These studies narrowed down the modifier regions to two sharp peaks and a third broader region between markers DXMit40 and DXMit116 [26]. The sharp peak showing strongest association with hypermuscularity is close to DXMit83 (27.99 cM), the second peak is bordered by DXMit130 (55.45 cM) and DXMit37 (59.10 cM) [26]. A survey of the genes located in the region of the peak at DXMit83 (27.99 cM) identified Usp26 (28.31 cM) of ubiquitin specific peptidase 26 (UniProt: Q99MX1), a component of the ubiquitin-dependent protein degradation system as a possible candidate that may modify the expressivity of the Compact phenotype. Our survey of the second major modifier region bordered by DXMit130 and DXMit37 failed to identify genes that might contribute to hypermuscularity of Compact mice, but in the region between DXMit40 and DXMit116, close to DXMit116 (49.58 cM) we find Ube2dnl1 (49.40 cM) and Ube2dnl2 (49.40 cM), ubiquitin-conjugating enzyme E2D N-terminal like 1 and 2 (UniProt: A2AFH1, and A2AFH2) that might function as constituents of the ubiquitin-dependent protein degradation system.

It should be pointed out that our suggestion that the phenotypic consequences of the misfolding of *Mstn*^*cmpt-Dl1abc*^ promyostatin are modified by constituents of the Unfolded Protein Response system is reminiscent of the conclusions drawn from studies on neurodegenerative diseases caused by misfolding of mutant proteins. In the last decade the search for modifier genes affecting susceptibility to neurodegenerative protein folding diseases has led to the conclusion that many of them encode proteins involved in protein folding and protein degradation [29, 30]. For example, the search for modifier genes of prion diseases led to the identification of Hectd2, an E3 ubiquitin ligase [31] and the chaperone Hspa13 [32] that is also a candidate modifier gene for the *Mstn*^*cmpt-Dl1abc*^ mutation (see above).

## 5. Conclusion

In the present work we have shown that the *Mstn*^*cmpt-Dl1abc*^ mutation impairs the folding and secretion of promyostatin. Nevertheless, the mutant precursor may give rise to some mature myostatin explaining why homozygous Cmpt/Cmpt mice have a less pronounced hypermuscular phenotype than homozygous mice in which the myostatin gene is disrupted [11]. This difference between Compact and myostatin-null mice may also be reflected in the spectrum of genetic loci that affect muscle mass: in the case of Compact mice components of the protein quality control system may increase the penetrance of the *Mstn*^*cmpt-Dl1abc*^ mutation (through efficient elimination of mutant promyostatin), but these proteins are not expected to have an influence on myostatin activity in myostatin null-mice. Our survey of the genes present in the chromosomal regions showing strong association with hypermuscularity of Compact mice has identified numerous members of the protein quality control system that might play a role in the elimination of *Mstn*^*cmpt-Dl1abc*^ mutant promyostatin.

In harmony with the difference between Compact mice and myostatin-null mice, the spectra of modifier loci identified in myostatin-null mice [33, 34] are markedly different from those defined using Compact mice [12, 13, 25, 26]: in the case of myostatin-null mice twelve Quantitative Trait Loci were identified on mouse chromosomes 1,3, 6, 7 that significantly interacted with the myostatin genotype [34], whereas in the case of compact mice hypermuscularity was associated with several unrelated markers, markers on chromosomes 16 and X showing the strongest association [12,13].

## Acknowledgements

This work was supported by grant 108630 of the National Scientific Research Fund of Hungary (OTKA).

